# Rapamycin not Dietary Restriction improves resilience against pathogens: a meta-analysis

**DOI:** 10.1101/2022.10.06.511121

**Authors:** Eleanor J Phillips, Mirre J P Simons

**Affiliations:** School of Biosciences, University of Sheffield, Western Bank S10 2TN Sheffield, United Kingdom

## Abstract

Dietary Restriction (DR) and rapamycin both increase lifespan across a number of taxa. Despite this positive effect on lifespan and other aspects of health, reductions in some physiological functions have been reported for DR and rapamycin has been used as an immunosuppressant. Perhaps surprisingly, both interventions have been suggested to improve immune function and delay immunosenescence. The immune system is complex and consists of many components. Therefore, arguably, the most holistic measurement of immune function is survival from an acute pathogenic infection. We reanalysed published post-infection short-term survival data of mice (n=1223 from 23 studies comprising 46 effect sizes involving DR (n=17) and rapamycin treatment (n=29) and analysed these results using meta-analysis. Rapamycin treatment significantly increased post infection survival rate (lnHR=-0.72; CI=-1.17, -0.28; p=0.0015). In contrast, DR reduced post-infection survival (lnHR=0.80; CI=0.08, 1.52; p=0.03). Importantly, the overall effect size of rapamycin treatment was significantly lower (P<0.001) than the estimate from DR studies, suggesting opposite effects on immune function. Our results show that immunomodulation caused by rapamycin treatment is beneficial to the survival from acute infection. For DR our results are based on a smaller number of studies, but do warrant caution as they indicate possible immune costs of DR. Our quantitative synthesis suggests that the geroprotective effects of rapamycin extend to the immune system and warrants further clinical trials of rapamycin to boost immunity in humans.

## Introduction

Ageing is the progressive decline of function and increased risk of death. Many phenotypes are associated with ageing (López-Otín et al., 2013), including declining immune function (Chung et al., 2002; Gavazzi and Krause, 2002). Immunosenescence leads to the dysfunction of immune cells affecting both innate and adaptive immunity (Nikolich-Žugich and Messaoudi, 2005; Ritz and Gardner, 2006; Shaw et al., 2013; Yousefzadeh et al., 2021) and to higher levels of inflammation (Baylis et al., 2013). Ageing therefore reduces our ability to mount an effective immune response, leaving us more susceptible to infection (Aw et al., 2007; Gavazzi and Krause, 2002). More broadly immunosenescence is thought to underlie several pathologies that appear during ageing, including cancer (Foster et al., 2011), autoimmune disease (Ritz and Gardner, 2006), as well as ineffective clearance and accumulation of senescent cells (Goronzy and Weyand, 2019; Yousefzadeh et al., 2021). Immunosenescence thus provides an attractive explanation and potential therapeutic avenue for ageing.

Established treatments that extend lifespan in model organisms, most notably dietary restriction (DR) (Fontana et al., 2010; Katewa and Kapahi, 2010) and mTOR suppression (Garratt et al., 2016; Johnson et al., 2013), might do so because they mitigate immunosenescence. The pro-longevity mechanisms of DR have been hypothesised to include mTOR suppression (Cox and Mattison, 2009; Green et al., 2022), but direct evidence for this hypothesis is scarce (Bjedov et al., 2010; Garratt et al., 2016; Miller et al., 2014; Unnikrishnan et al., 2020). Whether DR and mTOR suppression promote a healthier immune system and whether they do so through shared mechanisms is currently unclear. There are reports of beneficial effects of both of these pro-longevity interventions on immune function, yet there is also evidence to the contrary (Jolly, 2004; Mannick et al., 2014; Saunders et al., 2001). In addition, rapamycin (inhibiting mTOR) has been used as an immunosuppressant (Saunders et al., 2001) and a loss of immune defence is a hypothesised cost of DR (Speakman and Mitchell, 2011).

When measurements of the composition of the immune system are taken as proxies for immune health, extrapolation to overall organismal health is difficult. An additional complication is that such proxies are often studied under controlled, pathogen free, conditions (Camell et al., 2021; Goldberg et al., 2015). In comparison, acute survival to pathogens has received less attention, but provides a strong experimental and potentially translational paradigm to study the effects of DR and rapamycin. Pathogen infection is a pervasive problem that intensifies with age (Castle, 2000; Gavazzi and Krause, 2002). Treatments that enhance the effectiveness of the immune system to overcome infection are thus highly relevant. Conversely, should pro-longevity treatments simultaneously reduce the capacity to fight-off infection, the beneficial impact of DR and rapamycin on healthspan could be negated by reduced survival following naturally occurring infections (Johnson et al., 2013). We conducted a meta-analysis on studies in mice and found that survival after pathogen exposure was reduced by DR but improved with rapamycin.

## Results

DR had a significant negative effect on survival following pathogen exposure (Figure 1, lnHR=0.80; CI=0.08, 1.52; p=0.03). There was a large proportion of relative heterogeneity (I^2^=0.68; Q-test df=16, p<0.01). The small sample size of (seven) studies and variation in the recorded moderators were too small to perform any meaningful moderator analysis. This together with heterogeneity between studies and interdependency of effect sizes from the same study and using the same controls reduces the overall confidence in this result. It is unlikely however that variation between studies was due to mouse genotype or degree of DR, as all studies used the common inbred mouse strain, C57BL/6 and DR of 40% (Table S1). However, the only study to find a significant positive effect of DR (Mejia et al., 2015) used a parasitic model of infection and was the only study to use females. No publication bias was detected using a rank correlation (Kendall’s τ_b_=-0.25; p=0.18, Figure S3).

**Figure 1.**
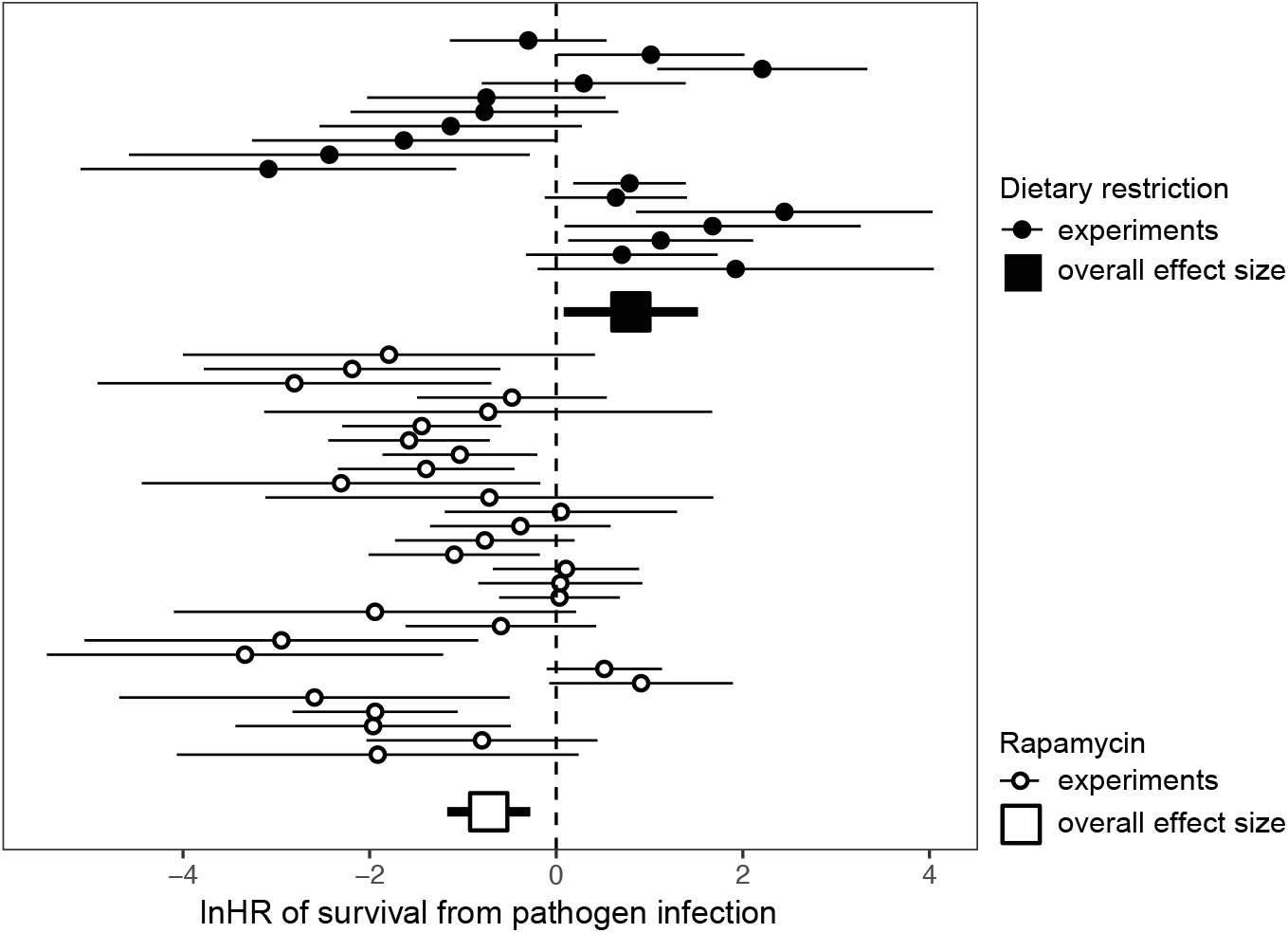
Forest plot of hazard ratio estimates (circles) for DR and rapamycin post-infection survival curve pairings (n=46) from Cox proportional hazard models. Squares indicate overall effect sizes as determined using meta-analysis controlling interdependence of study and shared controls. Whiskers indicate 95% CIs.

Rapamycin treatment improved survival of mice exposed to pathogens (lnHR=-0.72; CI=-1.17, -0.28; p=0.0015). Strikingly, when both interventions were analysed together, with treatment type as moderator, rapamycin treated mice had significantly better survival than those treated with DR (estimate=-1.50; CI: -2.33, -0.68; p < 0.001). There was large relative heterogeneity (I^2^=0.67; Q=84, df=28, p<0.01). To perhaps explain some of this heterogeneity we tested a number of possible moderators. We found no significant contribution from mouse genotype (Q_M_=2.78, df=4, p=0.60; or when testing BL6 against other: Q_M_=0.63, df=1; p=0.43), inoculation method (Q_M_=0.76, df=2; p=0.69), or pathogen type (Q_M_=1.25, df=3, p=0.74). The effect of sex could not be evaluated as information was not provided or was female (see Table S2). There was a trend that secondary infection (Q_M_=3.49, df=1, p=0.06) showed a stronger effect of rapamycin (−0.81; CI=-1.65, 0.04). A rank test of funnel plot asymmetry revealed no evidence for publication bias (Kendall’s τ_b_=0.23; p=0.09; Figure S4).

## Discussion

Through meta-analysis we found that rapamycin treatment but not DR significantly increased survival of mice exposed to pathogens. The pooled results of the limited number of studies suggest that DR does not improve immunity to infection and could even worsen the response. Studies on the impacts of rapamycin on infected mice have been inconclusive when comparing individual studies (Canivet et al., 2015; Huang et al., 2017). Contrary to DR, however, our meta-analysis revealed that rapamycin protected against pathogenic infection. This disparity between DR and rapamycin supports previous suggestions, that these two anti-ageing treatments operate though largely distinct mechanisms (Birkisdóttir et al., 2021; Garratt et al., 2016; Miller et al., 2014; Unnikrishnan et al., 2020).

A common interpretation is that DR benefits immune function by keeping it ‘younger for longer’ (Messaoudi et al., 2008; Pae et al., 2011). For instance, by protecting T-lymphocytes from oxidative damage (González et al., 2012), altering specific lymphocyte populations (Abe et al., 2001) and delaying thymic maturation (Chacón et al., 2002). However, our meta-analysis suggests that this ‘youthful’ immune system does not translate into a more potent response to pathogens. Perhaps aspects of innate immunity are compromised under DR. A reduced level of IL-6 (Sun et al., 2001) and reduced number and cytotoxicity of NK cells (Clinthorne et al., 2010) under DR were associated with reduced survival of mice upon infection. While DR decreases effectiveness of NK cell-based immunity, arguably regulated by leptin (Clinthorne et al., 2013, 2010; Naylor and Petri, 2016), this could also prevent a hyperimmune response enhancing survival. Similarly, a reduction in leptin production under DR was shown to be responsible for enhanced survival from cerebral malaria and these effects were mediated through reduced mTORC1 activity in T cells (Mejia et al., 2015).

Several mechanisms could explain why rapamycin increases resilience against pathogen infection. Immunosuppressive properties of rapamycin could prevent the activation of an overzealous immune response (Canivet et al., 2015; Kalil and Thomas, 2019). A more effective immune response could stem from elevated numbers of T regulatory (Treg) cells seen after rapamycin treatment (Canivet et al., 2015; Goldberg et al., 2014). Treg cells cause immune suppression to maintain homeostasis, for example reducing cytokine production which in turn ameliorates tissue damage (Liu et al., 2016). Rapamycin may also improve immune memory (Chen et al., 2009; Keating et al., 2013; Liepkalns et al., 2016), possibly fitting with the trend that secondary infections showed a stronger response to treatment. Rapamycin’s ability to reduce the debilitating effects of ageing on a systemic level could directly or indirectly benefit the immune system (Bischof et al., 2021). It remains to be determined to what degree the life-extending effects of rapamycin are due to its modulation of the immune system. Although lifespan extension by rapamycin in mice lacking T and B lymphocytes (RAG2^-/-^) without a rescue from an immune challenge (Hurez et al., 2015), suggests immunomodulation is not exclusively responsible for rapamycin’s anti-ageing effects. Outside the protected lab environment, however, infection and repeated exposure to pathogens could be strongly determinative of healthy ageing and lifespan. In this context, rapamycin has a strong immediate potential to benefit humans (Bischof et al., 2021).

For the studies included in our meta-analysis the duration and timing of treatment and age at pathogen exposure was so heterogeneous that we were unable to assess it (Table S2). Notably, in one study, short term rapamycin treatment was more successful in improving post-infection survival than long term treatment (Hinojosa et al., 2012). When comparing rapamycin to DR treatment we note that the majority of the DR studies initiated treatment well in advance of infection, whereas treatment with rapamycin was more brief. In fact, the one study that started DR on the day of infection was also the only study to find a significant benefit to survival (Mejia et al., 2015). Timing and scheduling of rapamycin treatment can have unpredictable effects and could depend on age. Transient rapamycin treatment (Juricic et al., 2022) and mTor knockdown (Simons et al., 2019) in early adult life extend lifespan in flies. Similarly, rapamycin during development (Shindyapina et al., 2022) and a short bout of treatment at middle-age (Bitto et al., 2016) extend lifespan in mice. Determining which rapamycin schedule is most beneficial to the ageing human will be key. It is encouraging however that short term rapamycin treatment in model organisms has benefits on both lifespan and on immune responses to pathogens, as we determined here through meta-analysis, paving the way for future human studies.

## Methods

### Literature Research

Scopus and Google scholar were the two primary databases used to collect results for search terms relating to both Dietary Restriction and Rapamycin. Additional sources were also found by searching the reference sections of salient papers [Denoted as ‘Other Sources’ in the PRISMA report – Figure S1]. As part of standard meta-analytic protocol (Shamseer et al., 2015) the PICO (Population, Intervention, Comparison, Outcome) framework was used to establish the specific research questions of the meta-analysis for both Rapamycin (How rapamycin impacts the immune response of non-mutant mice compared to mice treated with Placebo Vehicle Injection) and DR treatment (How DR impacts the immune response of non-mutant mice compared to mice fed ad libitum). From our initial literature research, we established that post infection survival is a common and relevant metric used. Although DR and rapamycin experiments have been conducted on species from a range of taxa, the most extensively studied and well controlled subject group were laboratory mice. Given this, we focussed the meta-analysis on studies on mice that measured short-term survival following pathogen exposure.

### Inclusion Criteria

General inclusion criteria: (i) The experiment contained a control group and a group under DR or treated with rapamycin. (ii) The study included survival data in the form of a Kaplan-Meier plot, or provided original/raw survival data. (iii) Studies that used mouse strains that were selected or genetically modified in a way that would prompt an abnormal response were excluded. For instance, p53 deficient mice were excluded as they exhibit accelerated immune ageing (Ohkusu-Tsukada et al., 1999). (iv) There were no restrictions on the age or sex, but this information was collected for potential use in moderator analysis. (v) Survival data from the experiment could be in response to primary pathogen exposure or secondary exposure to the same or similar pathogen. For instance, in a study by Keating and colleagues (2013). (vi) The studies chosen were restricted to those which used microparasites as the pathogen for their immune challenge. (vii) There were no restrictions on the date papers were published. (viii) Studies with insufficient or unclear data were excluded (e.g., studies that did not include sample size, or only survival data as an overall percentage rather than a Kaplan-Meier plot. One study such, by Huang and colleagues (2017), was due to a culmination of insufficient detail (rapamycin dose and mouse sex were not stated), a lack of independent controls and small sample size. Treatment specific inclusion criteria: For DR experiments: (i) Restrict overall food intake as opposed to restricting a specific macro or micronutrient. (ii) There was no limit on duration of DR prior to infection. (iii) Studies with DR conditions of 40-60% *ad libitum* to represent moderate restriction. For rapamycin experiments: (i) The experiment could use rapamycin at any dosage but not in conjunction with another drug. (ii) There was also no restriction on duration of rapamycin treatment, but this information was also recorded.

### Search Methodology

The following key terms were entered into the chosen databases, the searches were modified to fit the format of an advanced search in each database. Scopus: 1. (“Dietary Restriction” OR “Undernutrition”) AND ((infection OR influenza)) AND (mice) AND NOT (review) returned 64 hits. 2. “Rapamycin” AND (infection OR influenza) AND (mice) AND NOT (review) returned 853 hits. Google Scholar: 1. (Dietary Restriction OR DR) AND (immune challenge OR infection) AND (mice OR Mouse) returned ∼68,100 hits. 2. [Dietary Restriction] AND (infection OR immune response) AND [mice] AND “research paper” returned ∼162,000 hits. 3. “Dietary Restriction” AND (infection OR influenza) AND [mice] AND -review returned ∼603 hits. Note, alternative names for/forms of rapamycin also queried but these did not return any additional studies. Papers were assessed and selected manually following our inclusion and exclusion criteria and subsequently using the PRISMA guide (Figure S1). All literature searches were conducted by EP. A secondary non-structured search was conducted by MJPS as this can yield additional suitable literature. Later cross-referencing with the structured search yielded five additional suitable studies for the meta-analysis (Figure S1).

### Data Extraction and Re-Analysis

Raw survival times were extracted using image analysis of published Kaplan-Meier survival curves. These analyses were performed using the WebPlotDigitizer analysis software. This software uses labelled axes from the published survival curve to then measure the location of points on each survival curve (Garratt et al., 2016; Swindell, 2017). The extracted data was re-analysed using Cox Proportional Hazards to assess the relationship between post infection survival probability and DR or rapamycin treatment (R package: survival; function: coxph (Therneau et al., 2000)). Individuals still alive at follow up were right-hand censored. No individuals were censored in these studies during the experiment. The effect size estimates and Kaplan-Meier survival curves generated from this analysis were compared to those in the original publications to confirm that data had been extracted accurately and the direction of the effect corresponded to those reported in the original published work. We extracted pathogen type, infection method, sex and mouse genotype to be used in possible moderator analysis (Table S2). To include as many pertinent studies as possible, a range of pathogens were included, and pathogen type was extracted as a moderator. Longevity induced by rapamycin treatment has been shown to be differentially affected by sex, with greater lifespan increase in female mice than male mice at a verity of doses (Miller et al., 2014). Genotype has also been shown to impact lifespan of mice treated with both DR (Swindell, 2012) and rapamycin (Swindell, 2017). Additionally, there is evidence that the most common mouse models used in relevant studies, BALB/c and C57BL/6, exhibit distinctive immune responses when exposed to bacterial infection (Fornefett et al., 2018).

### Meta-Analysis

Effect sizes, expressed as log hazard ratios from each study were then analysed using a random effects multilevel meta-analysis model (R package: metafor; function: rma.mv (Viechtbauer, 2010). Standard errors from the Cox proportional hazard models provided the weighting of each effect size in the analysis (the inverse of s.e. squared). As several effect sizes estimated used the same control group, we accounted for this shared variance by including a covariance matrix (Garratt et al., 2016) calculated using ‘vcalc’ in metafor, using a correlation of 0.5 between effect sizes of shared controls. Multilevel meta-analysis allows the inclusion of random effects and we included study as a random intercept for the multiple experiments from the same study. Where possible, post hoc subgroup analysis was performed to assess potential variables that may have contributed to heterogeneity. We only performed moderator analysis if the moderator could be objectively coded as a continuous variable or a factor with enough replication within levels to be tested. We indicate in the text where this was not possible due to heterogeneity in reporting or low number of replications. Relative heterogeneity was assessed using a multilevel version of I^2^ (Nakagawa and Santos, 2012) and we also report Q tests. Publication bias within the meta-analysis was assessed visually using funnel plots (Figures S3 and S4) and statistically using a rank correlation test for funnel asymmetry using Kendall rank correlations.

## Acknowledgements

Funding from Sir Henry Dale Fellowship (Wellcome and Royal Society; 216405/Z/19/Z) and an Academy of Medical Sciences Springboard Award (the Wellcome Trust, the Government Department of Business, Energy and Industrial Strategy (BEIS), the British Heart Foundation and Diabetes UK; SBF004\1085).

## Supplementary material

**Figure S1:**
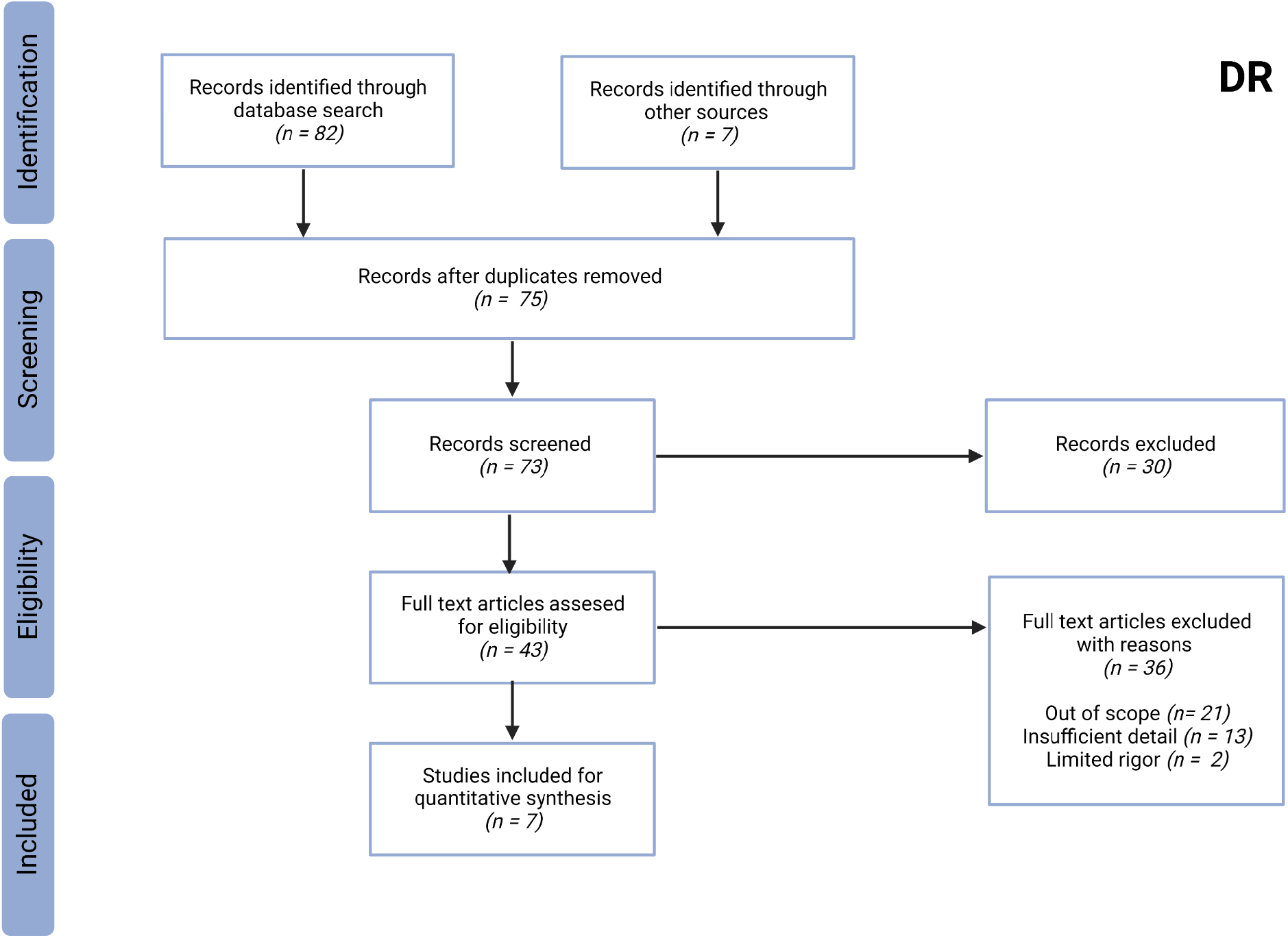
PRISMA flow chart describing search strategy for appropriate Dietary Restriction studies to be used in the present meta-analysis.

**Figure S2:**
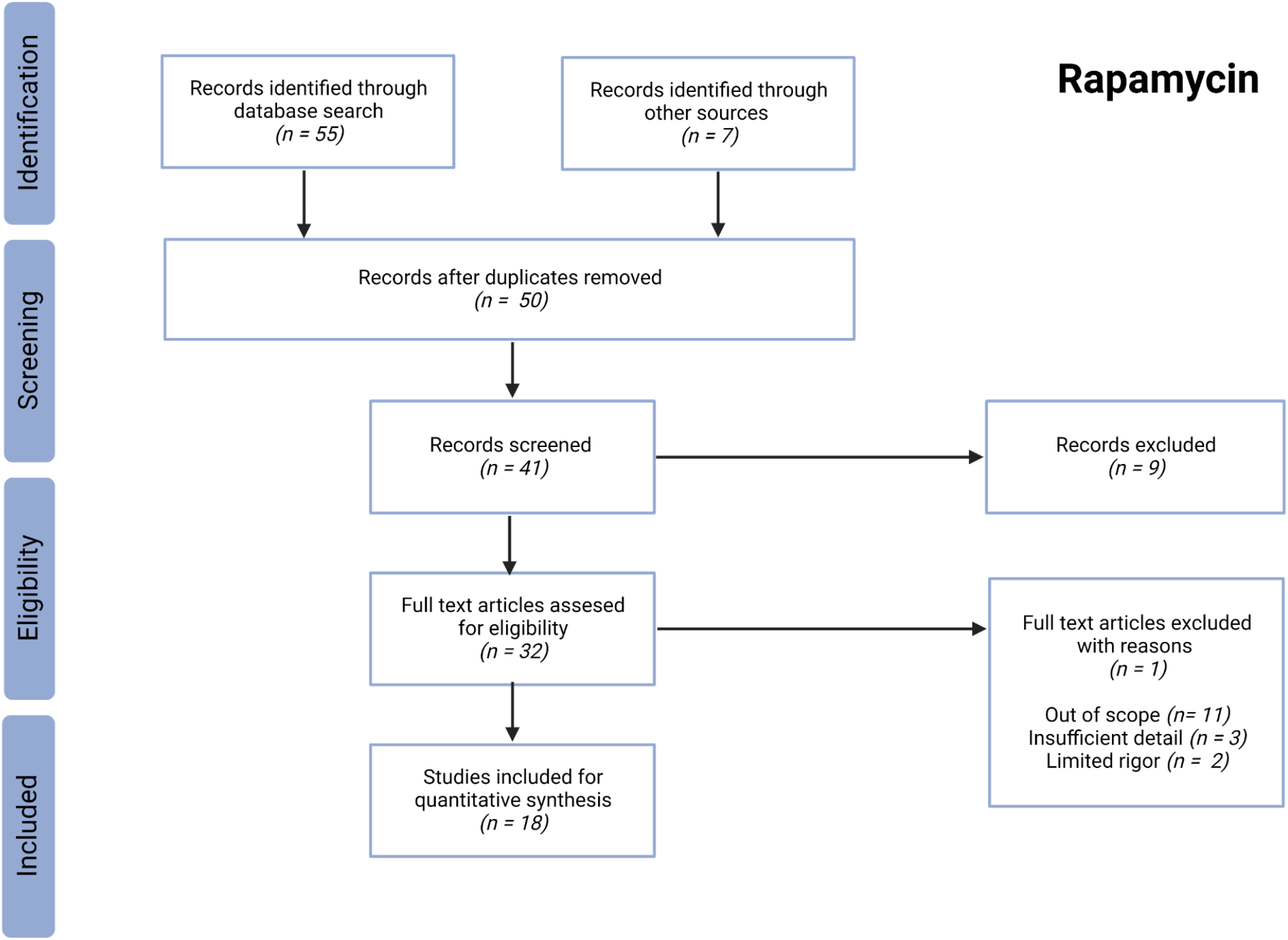
PRISMA flow chart describing search strategy for appropriate rapamycin studies to be used in the present meta-analysis.

**Table S1:**
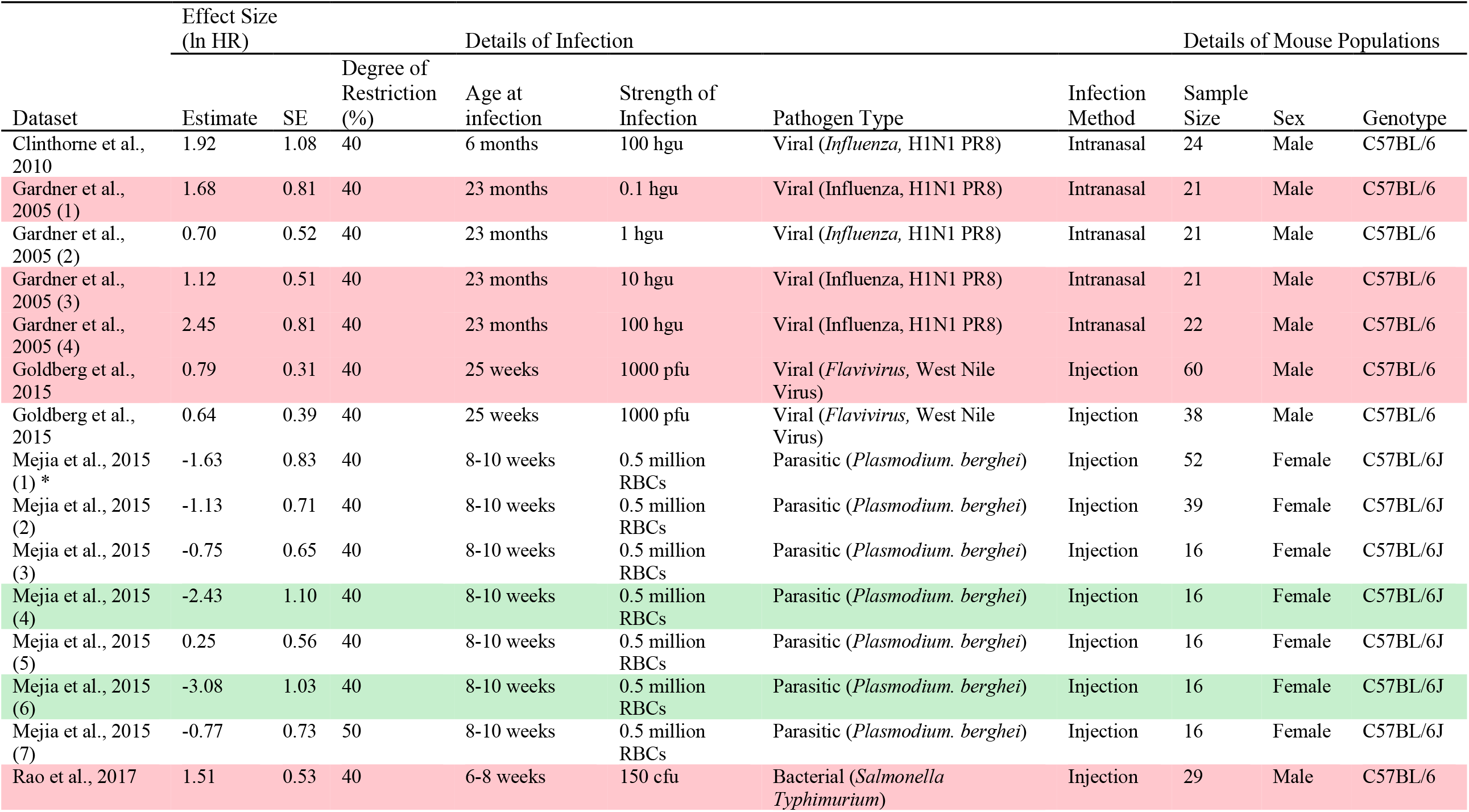
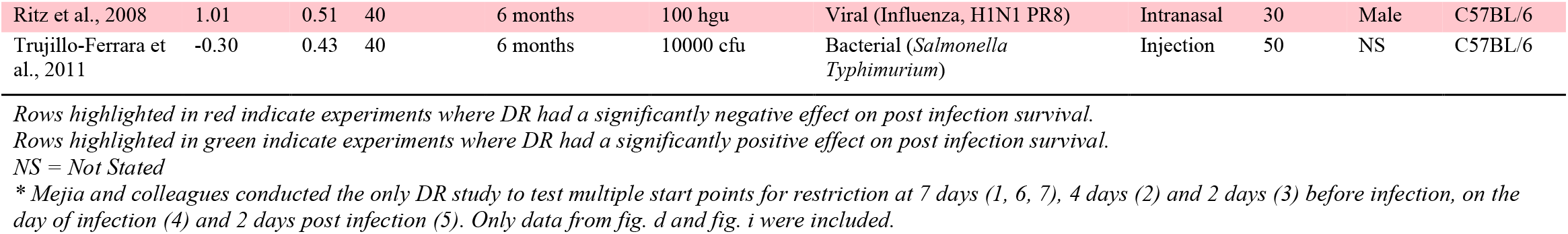
Additional details of Dietary Restriction studies used in the meta-analysis, including effect sizes and details of DR treatments, infections and mouse populations used in each experiment.

**Table S2:** Additional details of rapamycin studies used in the meta-analysis, including effect sizes and details of DR treatments, infections and mouse populations used in each experiment.

| Dataset | Effect Size<br>(ln HR) |  | Details of Rapamycin Treatment |  | Details of Infection |  |  |  |  | Details of Mouse Population |  |  |
| --- | --- | --- | --- | --- | --- | --- | --- | --- | --- | --- | --- | --- |
|  | Estimate | SE | Rapamycin Dose* | Frequency of Treatment | Age at Infection | Priming Infection | Pathogen Type | Infection Strength | Infection Method | Sample Size | Sex | Genotype |
| Bell et al., 2017 | -1.91 | 1.10 | 10 mg/kg daily | 1 hour before infection to 10 days post infection | 6-8 weeks | No | Viral (Rift Valley Fever Virus) | 150 pfu | Injection | 20 | Female | BALB/c |
| Bell et al., 2017 | -0.80 | 0.63 | 10 mg/kg daily | 1 hour before infection to 14 days post infection | 6-8 weeks | No | Viral (Rift Valley Fever Virus) | 1500 pfu | Injection | 20 | Female | BALB/c |
| Canivet et al., 2015 | -1.96 | 0.75 | 10 mg/kg daily | Days 4 to 13 post infection | 4-5 weeks | No | Viral (Herpes Simplex Virus) | $1.5 \times 10^3$ pfu | Intranasal | 28 | Female | BALB/c |
| Canivet et al., 2015 | -1.94 | 0.45 | 10 mg/kg daily | Days 4 to 13 post infection | 4-5 weeks | No | Viral (Herpes Simplex Virus) | $1.5 \times 10^3$ pfu | Intranasal | 45 | Female | BALB/c |
| Chen et al., 2009 | -2.59 | 1.07 | 4mg/kg every 2 days | 8 weeks before infection | 22-24 months | Yes | Viral (*Influenza* H1N1 PR8) | 400 hau | Intranasal | 24 | NS | C57BL/6 |
| Goldberg et al., 2014 | 0.91 | 0.50 | 75µg/kg daily | 2 days before infection to 7 days post infection | 16-18 months | No | Viral (*Flavivirus*, West Nile Virus) | $10^3$ pfu | Injection | 42 | NS | C57BL/6 |
| Goldberg et al., 2015 | 0.52 | 0.32 | 75µg/kg daily | 2 months before infection | 16-18 months | No | Viral (*Flavivirus*, West Nile Virus) | 400 hau | Intranasal | 57 | NS | C57BL/6 |
| Gordon et al., 2015 | -3.34 | 1.08 | 1mg/kg daily | 1 day post infection (until experiment end) | 7-10 weeks | No | Parasitic (*Plasmodium berghei*) | $1 \times 10^6$ RBCs | Injection | 19 | Female | C57BL/6 |
| Gordon et al., 2015 | -2.95 | 1.08 | 1mg/kg daily | 4 days post infection (until experiment end) | 7-10 weeks | No | Parasitic (*Plasmodium berghei*) | $1 \times 10^6$ RBCs | Injection | 18 | Female | C57BL/6 |
| Gordon et al., 2015 | -0.59 | 0.52 | 1mg/kg daily | 5 days post infection (until experiment end) | 7-10 weeks | No | Parasitic (*Plasmodium berghei*) | $1 \times 10^6$ RBCs | Injection | 18 | Female | C57BL/6 |
| Gust et al., 2011 | -1.94 | 1.10 | 75µg/kg daily | 1 day before priming infection until secondary infection (28 days) | NS | Yes | Viral (*Influenza* H5N1) | $1 \times 10^8$ EID<sub>50</sub> | Injection | 20 | Female | C57BL/6 |
| Harrison et al., 2014 | 0.04 | 0.33 | NS | 6 weeks prior to infection and withdrawn before infection | 24.5 months | No | Bacterial (*Mycobacterium tuberculosis* (GP)) | NS | Inhalation | 50 | Female | HET3 |
| Heydarabadi et al., 2020 | 0.04 | 0.45 | 40µg/µl daily | NS | NS | No | Viral (*Rhabdoviridae*, RABV) | NS | Injection | 20 | NS | NMRI |
| High and Washburn, 1996 | 0.10 | 0.40 | 10mg/kg | 1 day before infection to 14 days post infection | NS | No | Fungal (*Aspergillus fumigatus*) | $7.5 \times 10^6$ conidia | Injection | 40 | NS | CD-1 |
| Hinojosa et al., 2012 | -1.09 | 0.47 | 2.2mg/kg daily | 17 weeks before infection | 24 months | No | Bacterial (*Streptococcus pneumoniae* (GP)) | $1 \times 10^3$ cfu | Inhalation | 25 | Both | C57BL/6 |
| Hinojosa et al., 2012 | -0.77 | 0.49 | 2.2mg/kg daily | 86 weeks before infection | 24 months | No | Bacterial (*Streptococcus pneumoniae* (GP)) | $1 \times 10^3$ cfu | Inhalation | 29 | Both | C57BL/6 |
| Jai et al., 2018 | 0.05 | 0.64 | 600µg/kg on day 1 then 300µg/kg daily | 2 hours post infection (until experiment end) | 6-8 weeks | No | Viral (*Influenza* H1N1 pdm09) | $10^2$ TCID<sub>50</sub> | Intranasal | 16 | Female | BALB/c |
| Jai et al., 2018 | -0.38 | 0.49 | 600µg/kg on day 1 then 300µg/kg daily | 2 days post infection (until experiment end) | 6-8 weeks | No | Viral (*Influenza* H1N1 pdm09) | $10^2$ TCID<sub>50</sub> | Intranasal | 32 | Female | BALB/c |
| Junkins et al., 2013 | -0.72 | 1.22 | 10mg/kg daily | 3 days before infection (until experiment end) | 8-10 weeks | No | Bacterial (*Pseudomonas aeruginosa* (GN)) | $10^9$ cfu | Intranasal | 30 | NS | C57BL/6 |
| Keating et al., 2013 | -2.31 | 1.09 | 75µg/kg daily | 1 day before primer infection | 8-10 weeks | No | Viral (*Influenza* A/HK/x31) | $1 \times 10^8$ EID<sub>50</sub> | Injection | 18 | Female | C57BL/6J |
| Keating et al., 2013 | -1.39 | 0.48 | 75µg/kg daily | 1 day before priming infection to 28 days post priming infection | 12-14 weeks | Yes | Viral (*Influenza* H5N1) | $4.5 \times 10^5$ EID<sub>50</sub> | Intranasal | 32 | Female | C57BL/6J |
| Keating et al., 2013 | -1.03 | 0.42 | 75µg/kg daily | 1 day before priming infection to 28 days post priming infection | 12-14 weeks | Yes | Viral (*Influenza* PR8) | $4.5 \times 10^5$ EID<sub>50</sub> | Intranasal | 36 | Female | C57BL/6J |
| Kim et al., 2020 | -1.58 | 0.44 | 150 µg | 3 and 5 hours after priming and 18 hours after secondary infection | 10 weeks | Yes | Fungal (*Candida albicans*) | $2 \times 10^4$ cfu | Injection | 28 | Female | C57BL/6 |
| Kim et al., 2020 | -1.44 | 0.43 | 150 µg | 3 and 5 hours after priming and 18 hours after secondary infection | 10 weeks | Yes | Fungal (*Candida albicans*) | $2 \times 10^4$ cfu | Injection | 28 | Female | C57BL/6 |
| Liepkalns et al., 2016 | -0.47 | 0.52 | 1.5 µg daily | 3 days before infection (until experiment end) | 6 weeks | No | Viral (*Influenza* H1N1 PR8) | 1.5 LD<sub>50</sub> | Intranasal | 26 | NS | C57BL/6 |
| Liepkalns et al., 2016 | -0.73 | 1.22 | 12 µg daily | 3 days before infection (until experiment end) | 6 weeks | Yes | Viral (*Influenza* H1N1 PR8) | 1.5 LD<sub>50</sub> | Intranasal | 20 | NS | C57BL/7 |
| Mejia et al., 2015 | -2.81 | 1.08 | 1 mg/kg | Days 1 to 3 post infection | 8-10 weeks | No | Parasitic (*Plasmodium berghei*) | 0.5 million RBCs | Injection | 20 | Female | C57BL/6 |
| Mejia et al., 2015 | -2.19 | 0.81 | 5 mg/kg | Days 1 to 3 post infection | 8-10 weeks | No | Parasitic (*Plasmodium berghei*) | 0.5 million RBCs | Injection | 20 | Female | C57BL/6 |
| Moraschi et al., 2021 | -1.79 | 1.13 | 0.075 mg/kg daily | 34 days starting at priming infection | 8 weeks | Yes | Parasitic (*Trypanosoma cruzi*) | 150 blood trypomastigotes | Injection | 14 | Both | C57BL/6 |

*Rows highlighted in green indicate experiments where rapamycin had a significantly positive effect on post infection survival.*
*No experiment showed rapamycin to have a significantly negative effect on survival.*
*NS = Not Stated*
*\*Refers to body weight per mouse.*

**Figure S3:**
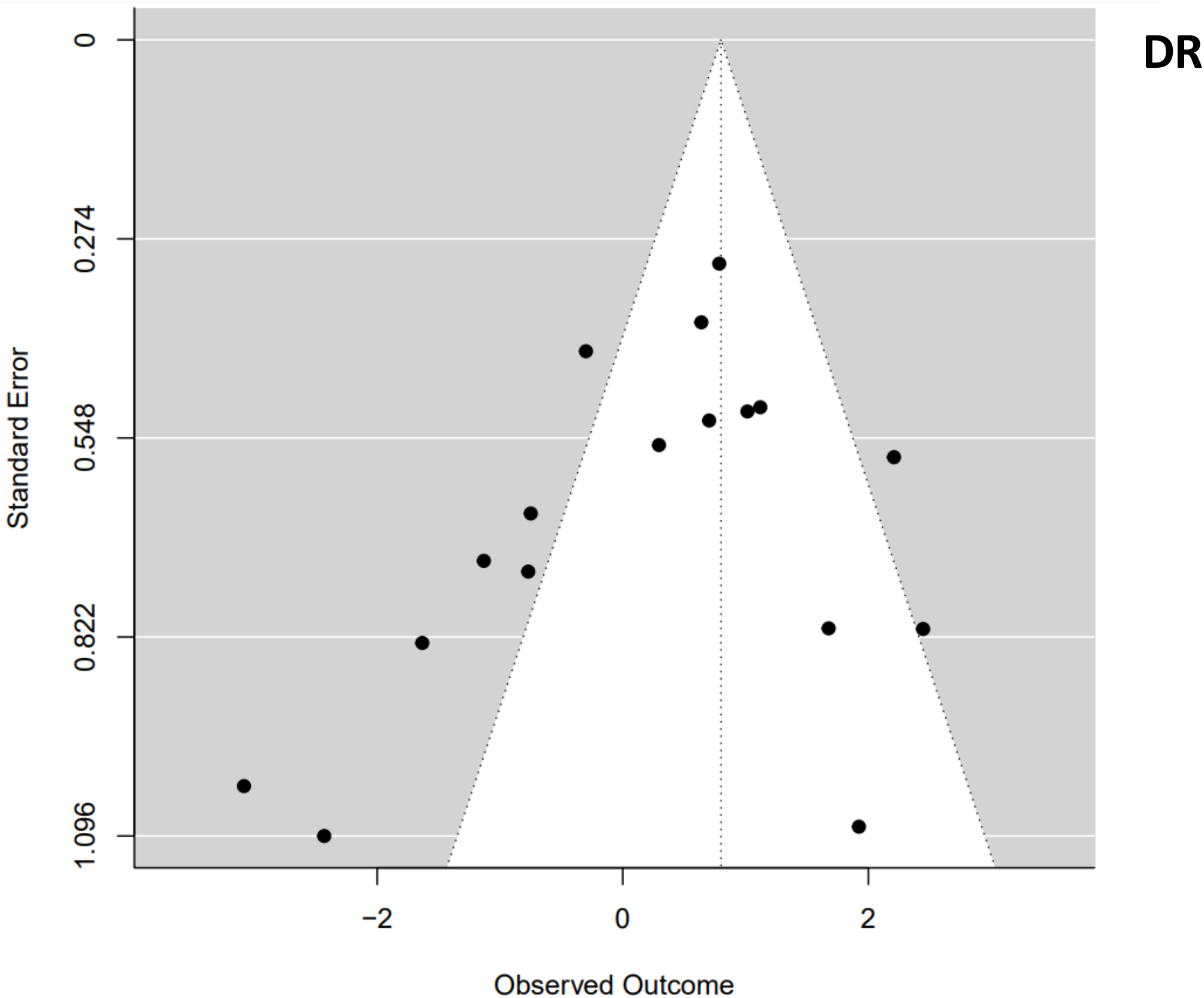
Funnel plot of the hazard ratio estimates of survival curve pairs taken from DR studies. The white triangle represents the 95% confidence interval that all experimental hazard ratio estimates would be expected to fall within if no publication bias is present (Kendall’s τ_b_ rank correlation test, p=0.18).

**Figure S4:**
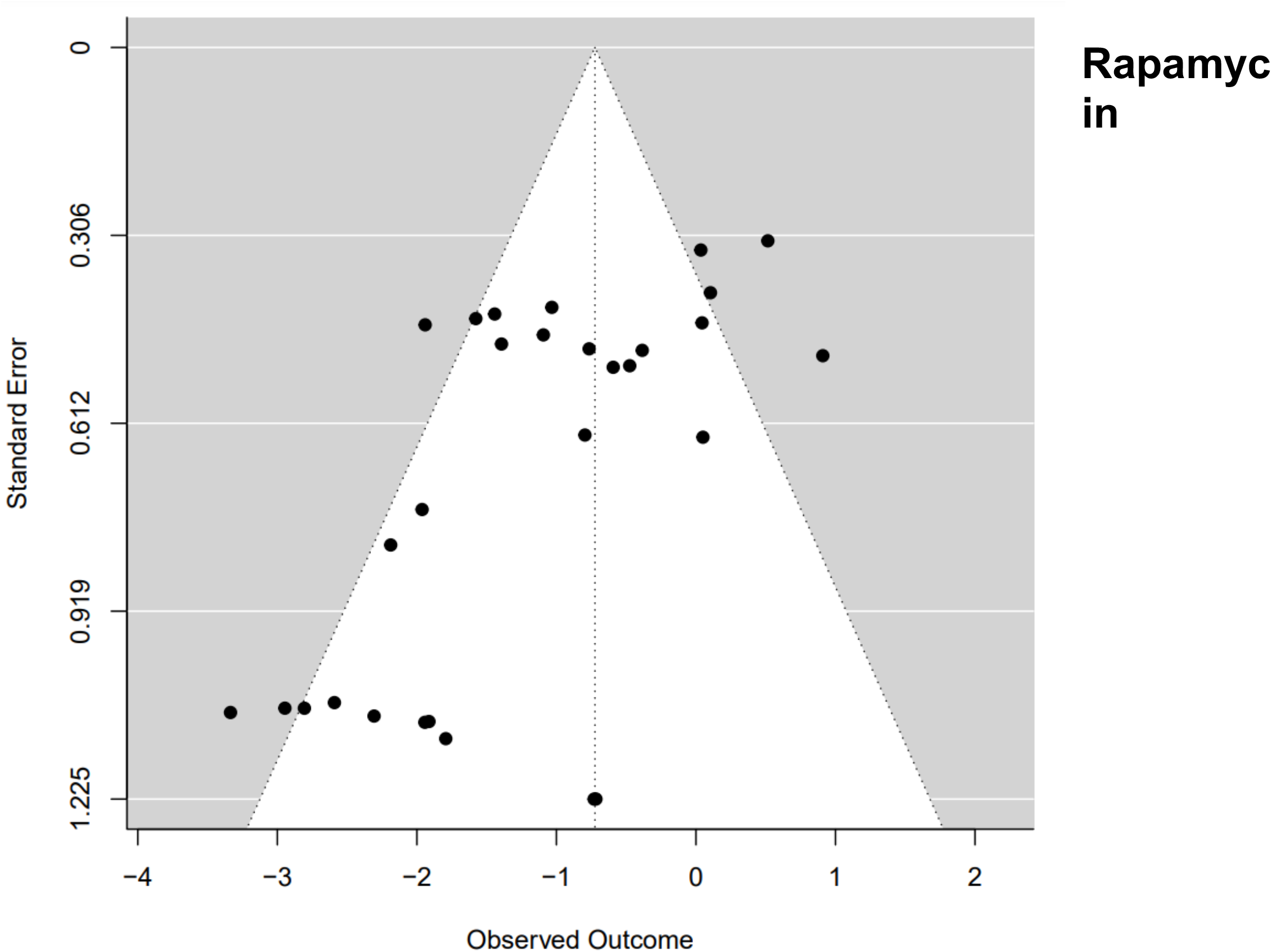
Funnel plot of the hazard ratio estimates of survival curve pairs taken from rapamycin studies. The white triangle represents the 95% confidence interval that all experimental hazard ratio estimates would be expected to fall within if no publication bias is present (Kendall’s τ_b_ rank correlation test, p=0.09)

